# Vapor-based Fixation of Pulmonary Tissue in its Physiological State: A Novel Approach to Histological Validation of Ultra High Resolution Phase Contrast CT in Human Sized Lungs

**DOI:** 10.1101/2024.08.26.609748

**Authors:** Christian Dullin, Johanna Reiser, Willi Wagner, Elena Longo, Marko Prašek, Adriano Contillo, Nicola Sodini, Diego Dreossi, Paola Confalonieri, Francesco Salton, Marco Confalonieri, Elisa Baratella, Maria Cova, Claudia Benke, Md Motiur Rahman Sagar, Lorenzo D’Amico, Jonas Albers, Angelika Svetlove, Tatiana Flisikowska, Krzysztof Flisikowski, Mark O. Wielpütz, Juergen Biederer, Hans-Ulrich Kauczor, Frauke Alves, Fabrizio Zanconati, Giuliana Tromba

## Abstract

Lung diseases continue to present a major burden on public health. Therefore, improving the process of diagnosis by the development of novel imaging techniques is of great importance. In this perspective, phase sensitive CT imaging techniques such as propagation based imaging (PBI) might play an important role as they allow increasing the spatial resolution at very low x-ray dose levels that are comparable to clinical CT. However, the development of such methods is not only hindered by technological problems but also by the lack of precise validation strategies. We adapted formaldehyde (FA) vapor fixation to demonstrate that fresh porcine lungs that have been investigated by PBI can be fixed in their physiological shape and studied by multi-scale microCT imaging as well as classical histology. In addition, we show that FA vapor fixed pig lungs can be scanned by PBI without visible deterioration of image quality compared to fresh tissue. This opens the possibility of fixing and storing, for instance, human lung tissue before performing a PBI experiment, which in turn allows to study pathological changes in human lungs without questioning the translate-ability of findings in pig lung. The setup can be used by any interested researchers.

## Introduction

Propagation based imaging (PBI) - the most commonly used x-ray phase contrast approach - has proven to provide strongly elevated soft-tissue contrast at very low dose rates, especially in pulmonary tissue (Kitchen et al., 2017). Therefore, PBI would be the ideal technology to be used in clinical CT imaging of the lung. However, as PBI requires a coherent x-ray source, long sample-to-detector distances and detectors with small pixel sizes, such an application will most likely be limited to synchrotron light sources in the near future. In this context, efforts are underway at the Italian and Australian light sources to implement the approach. A synchrotron PBI based lung CT will be performed as an additional local area examination - resembling a kind of virtual lung biopsy. Therefore, patients with a non-conclusive clinical CT or patients in which taking invasive lung biopsies is strongly contraindicated would benefit the most from such a new imaging method.

Prior to the construction of a clinical facility and in light of applying for ethical and technical authorization, such a method needs to be compared to clinical standards like conventional pathohistology for the examination of tissue specimens. The challenge is to perform imaging as well as the validation in conditions as close as possible to the already established way of clinical diagnosis. We already demonstrated that by the use of an anthropomorphic human chest phantom, fresh unfixed porcine lungs can be analyzed by both synchrotron PBI as well as clinical CT. The resulting images bear a striking resemblance to the patient’s CT scan. However, the porcine lungs used in the study were obtained from licensed slaughterhouses and were therefore free from obvious pathological changes. So far, we have simulated lung pathologies by injecting warm agarose gel mixed with iodine Albers et al. (2023). While the radiological pattern looked similar to the appearance of pulmonary disease in patients, histological validation of the imaging findings was not possible. Real pathologies and the usage of human lung tissue would therefore be greatly beneficial for optimizing and validating the method.

FOr safety reasons, human lung specimens, for instance from body donors or post lung transplantation, can not be used without fixation and in most of the cases do not represent a whole, undamaged lung. Thus, a fixation method for the lung tissue is needed and should support two scenarios: A) fixing a fresh porcine lung within the phantom after CT acquisition to maintain the same expansion state of the lung and allow for an easier comparison of subsequent histological analysis and B) fixing, for instance, human lung tissue within the phantom prior to CT acquisition to preserve the tissue for a following imaging session. Fixation by pouring Paraformaldehyde (PFA) into the lung is not ideal as the lack of air within the airways reduces the efficacy of PBI dramatically.

We present our findings using formaldehyde (FA) vapor fixation of pulmonary tissue in its physiological expansion state based on the method proposed by Weibel and Vidone (Weibel and Vidone, 1961) to compare PBI based virtual biopsy with clinical bronchoscopy and histology of lung biopsies.

## Results

### Validation problem in PBI based lung CT

In Fig.1a) a PBI scan of an unfixed porcine lung within the chest phantom is shown. The lung was obtained from a mini-pig that had been diagnosed and treated for pneumonia. Clearly, the highlighted area had an increased density within the CT scan and appeared to be pathologically altered. To validate and further analyse this finding, histology was needed. Once removed from the phantom, the lung collapses and the localization and identification of the same specific region was challenging (Fig.1b) and c)). In this specimen, the lung region was severely altered not only within the CT scan but also on a macroscopic level. FFPE-tissue blocks from the pathologically altered region (Fig.1d) turquoise) and from a supposedly healthy control region (Fig.1d) green) were generated. Fig.1e) and f) show an overview of a H&E stained slice from both of these blocks. The tissue that had supposedly been affected by pneumonia (Fig.1e)) appears more dense and its shape seems to resemble the shape in the PBI data set. Nevertheless, the lung tissue appears to be compressed and distorted. The overall quality of the histological images is not good and the deformation of the lung after being removed from the phantom rendered the correlation to the CT scans very challenging. To combat this, we developed a new fixation process which was validated by scanning the lungs in their fresh, unfixed state at negative pressure and subsequently without negative pressure after the vapor fixation.

**Figure 1.**
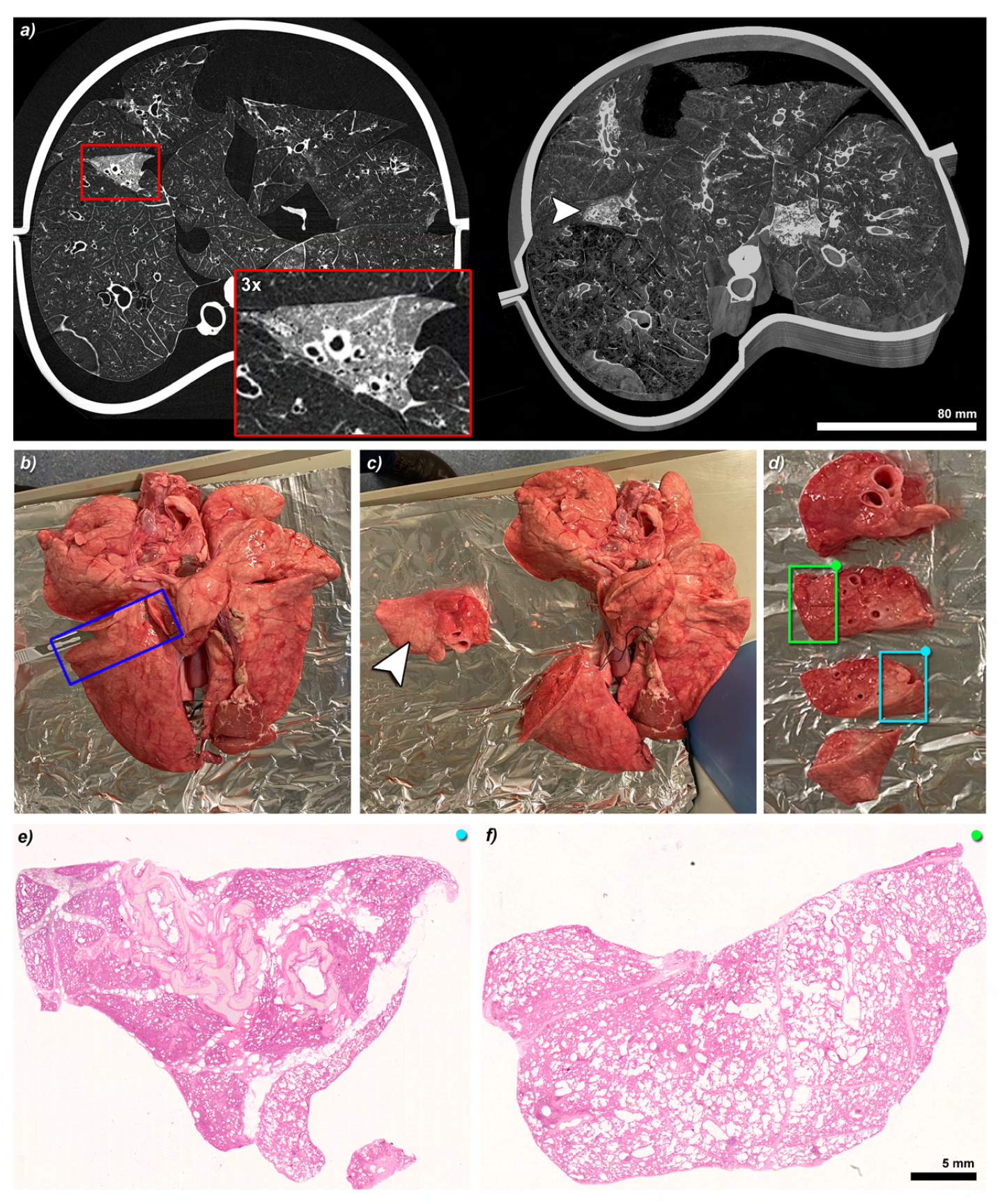
PBI imaging results of a lung retrieved from a pig after recovery from pneumonia and comparative histology. (**a**) shows a slice (left) and a 3D rendering (right) of a PBI scan of a porcine lung within the phantom. The pig had recovered from pneumonia. In the magnified region, pathological changes in one lung lobe can be seen. Scale bar 80 mm. (**b**) depicts the collapsed lung removed from the phantom. The blue rectangle indicates the region that was further processed. (**c**) shows that lung tissue was macroscopically more dense and pale in the targeted area (arrow head). (**d**) shows the approximately 1 cm thick lamella obtained from this piece of tissue. The turquoise and the green rectangle indicate the pathologically altered tissue and the supposedly healthy tissue that was later FA fixed, chemically dehydrated and embedded in paraffin. (**e**) and (**f**) show H&E stained sections of those regions. Scale Bar, 5 mm. Clearly, the pathological tissue in (**e**) was more dense and its shape had a strong resemblance to the shape of the dense structure identified in PBI (**a**, insert). Overall, the quality of the histological sections is rather poor. The lung tissue seems to be damaged, distorted and collapsed.

### FA vapor fixation setup

The essential condition for the high image quality in PBI is the huge difference in the refractive index between the lung tissue and the air filled bronchi. Therefore, a fixation process that would instill FA or PFA into the airways of the lung would cause a dramatic decrease of image contrast. To bypass this, we developed a FA vapor based fixation approach. In Fig.2a) the lung phantom is shown. This system consist of: i) an inner shell in which a lung can be placed with the trachea connecting to the outside, ii) an outer shell to mimic attenuation of chest wall and bones in patients, iii) a vacuum pump (on top) which realized a negative pressure between lung and phantom, resulting in a physiological passive inflation of the lung and iv) a motion system that allows simulation of breathing motions. Fig.2b) shows the concept presented by Weibel and Vidone (1961). In essence, FA is evaporated by boiling 10% formalin, resulting in the production of steam. This steam is then drawn into the lung via the trachea, due to the negative pressure created within the box housing the lung, which is generated by a flow pump. The red dashed part of the concept is already core functionality of the lung phantom, which therefore can be repurposed for the fixation process. Fig.2c) shows the final setup placed within a security working bench. The inner shell of the phantom (**N**) is equipped with a fresh porcine lung and mounted upright to exploit the bottom part as a reservoir for a potentially high amount of liquid. The phantom is connected to a vacuum pump (**P\***) which is keeping the lung inflated by sucking in air through the trachea/pipe. The trachea/pipe is connected via tubing’s with the FA evaporator (**A**), which in our case was a simple heating plate to boil 10% formalin (4% FA). The FA vapor is cooled by a water bath (**W**). On the pump side we used a three-way switch (**H**) to regulate the suction rate and a scavenger system made from two bottles (first filled with water) (**B**) to avoid letting FA vapor and liquid enter the pump (**P\***). Fig.2d) shows the appearance of the lung after approximately 4 hrs of fixation. The brownish color (**\***) indicates complete penetration of the FA, reaching the surface of the lung. In the area indicated in (**§**), the lung was attached to the inner surface of the phantom limiting the flow of FA vapor. Its reddish color demonstrates that this area was not yet fixed completely. After approximately 6 hrs, a complete fixation of the lung was achieved. To remove any additional fluid buildup, the bottom part of the phantom was slightly. Afterwards, the lung was scanned again with the PBI setup without applying negative pressure to the chest phantom.

**Figure 2.**
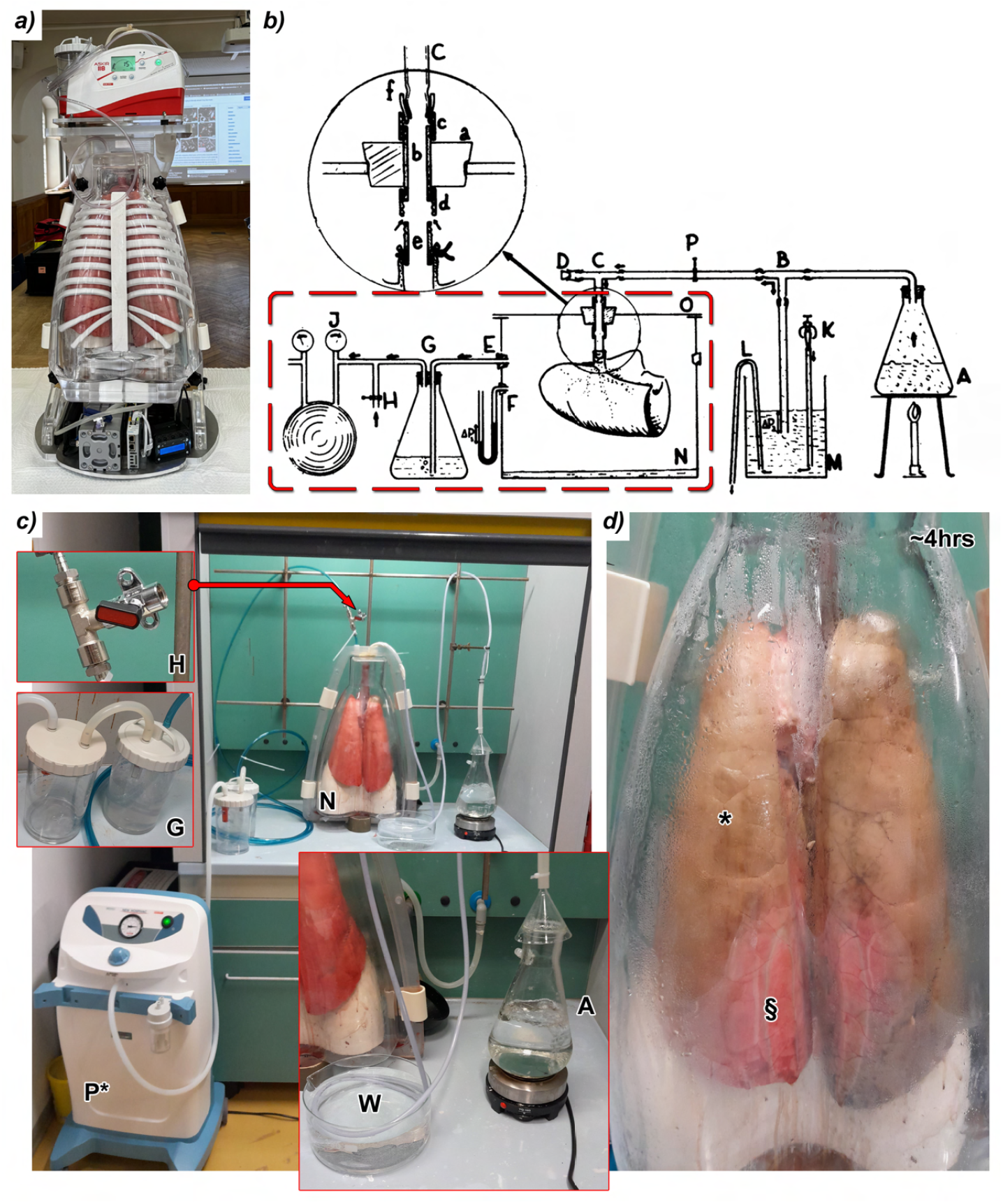
Concept for FA vapor fixation of the expanded lung. (**a**) shows the chest phantom with an inserted unfixed pig lung. The lung was kept passively inflated using negative pressure generated by the pump seen on top. (**b**) shows the vapor fixation concept presented by Weibel and Vidone (1961). FA vapor is generated by boiling FA **A**. The lung is placed in a box **N** in which a negative pressure can be generated by the pump **J**. The lung’s bronchial tree is connected to a pipe system **C** resulting in an influx of the FA vapor into the lung. The part outlined in red represents the existing functionality of the used lung phantom. (**c**) shows the vapor fixation setup placed in a security workbench. **N** depicts the inner part of the phantom containing the lung with the trachea fixed to a pipe. Negative pressure is applied by the flow pump **P\***. The valve **H** was used to adjust the pressure. **G** denotes a water filled scavenger to avoid FA vapor from entering the pump. **A** depicts the FA boiling system and **W** the water bath used to cool down the vapor before entering the lung. (**d**) shows the lung after 4 hrs of fixation. The main part of the surface of the lung (*) has turned brownish, indicating a full penetration of the FA. Some regions (§) attached to the inner surface of the phantom were not fixed entirely at this time, but after the whole process of 6 hrs.

### Comparison of PBI imaging of fresh and FA vapor fixed lungs

An important aspect was to validate that the FA vapor fixed tissue can be used for PBI imaging without a significant decrease of achievable image quality. This would facilitate the processing of human lung material, as it could be fixed in advance. Fig.3a) shows a 3D rendering of an unfixed mini-pig lung scanned within the lung phantom with PBI while using the negative pressure pump to keep the lung passively inflated. The data has been virtually cut open in order to more effectively illustrate the level of image quality that has been achieved. Fig.3b) shows the same lung scanned after 6 hrs of FA vapor fixation, which together with setup times resulted in 11.5 hrs difference between the two scans. For the scan of the fixed lung the pressure pump was not used. Apart of some minor deformations, the images Fig.3a) and b) demonstrate a high level of resemblance between the two conditions of the lung. Most importantly, the image quality of the PBI scans was not diminished after fixation. Roughly 4 weeks later, the FA vapor fixed lung which was stored in a sealed bucket at room temperature was sliced for further histological assessment. Fig.3c) shows that the slice microscopically shows the same details. Please note that the virtual cut in the CT data set shown in Fig.3a) and b) were retrospectively selected to match the appearance of Fig.3c). An approximately 5 mm thick layer of this lung lobe was further split into pieces of approximately 2 × 3 cm^2^, chemically dried and paraffin embedded. Fig.3d) demonstrates an ensemble of H&E stained sections of those blocks close to the surface of the tissue. Apart from the known deformations during the cutting process, the same structures seen in Fig.3 were found. The magnified views of the positions indicated by the looking glasses (Fig.3e) and f)) indicate a pristine tissue preservation.

**Figure 3.**
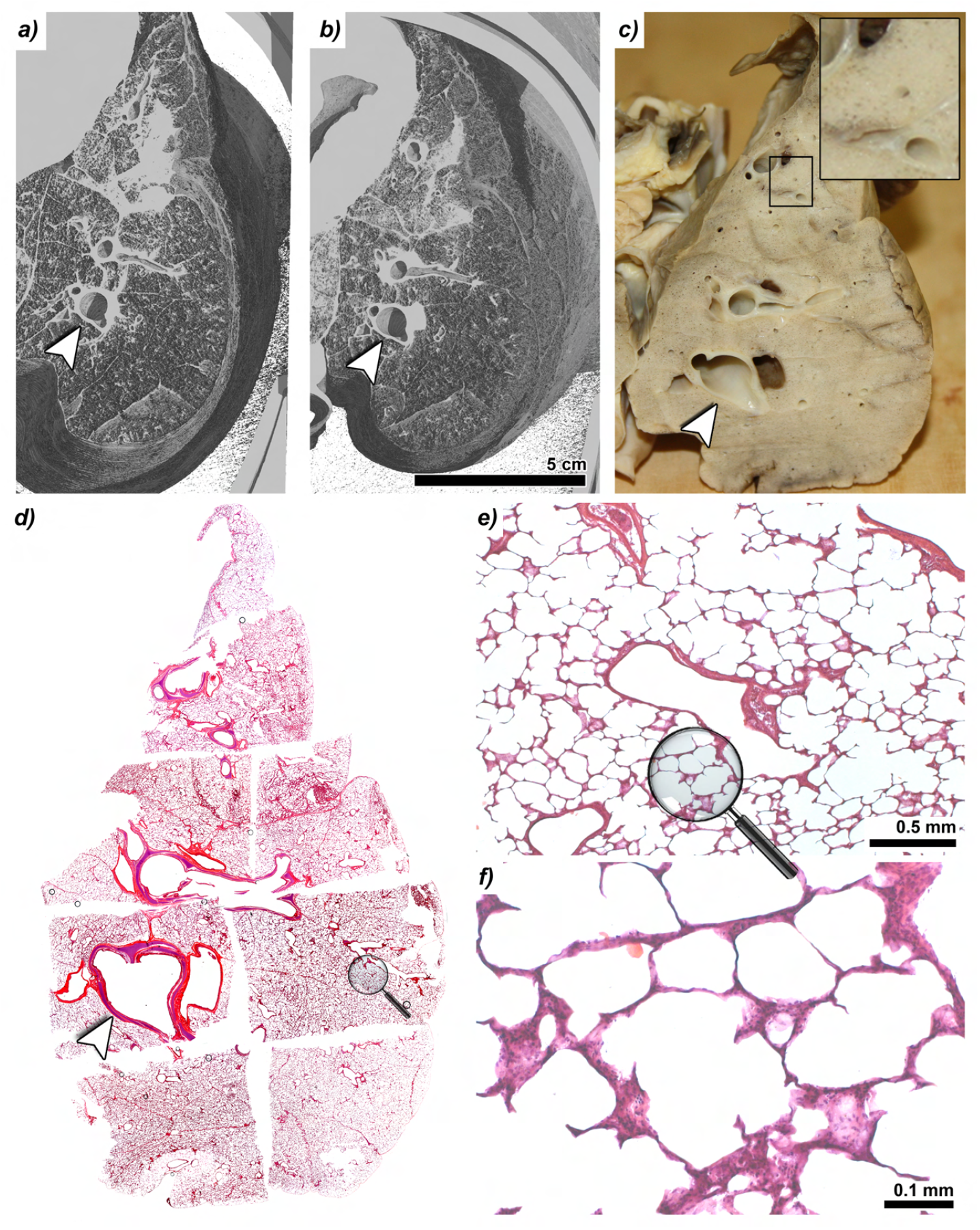
Comparison of fresh lung tissue vs. FA fixed tissue. (**a**) shows a 3D rendering of a fresh pig lung within the phantom. (**b**) shows the same lung after 6 hrs of FA vapor fixation (scan performed 11.5 hrs after the first one and without negative pressure during the scan). No apparent deterioration of the PBI image quality was found after FA vapor fixation. Scale bar 5 cm. (**c**) shows the same lung processed for histological examination approximately 4 weeks later. The detail view shows the successful preservation of the physiologically porous nature of the lung and the absence of liquid in the bronchi. Please note that since it was uncertain whether the heart would be completely fixed as well, it was removed prior to fixation. 9 FFPE-tissue blocks were obtained from the entire lung slice. (**d**) shows the assembled H&E stained superficial slices. (**e** and **f**) show magnified regions at the positions indicated by the looking glasses. Also at this magnifications a satisfactory preservation of the tissue was found.

### FA vapor fixation enables multi-scale imaging

The ability to fix the lung after the PBI scan in the same physiological shape enables multi-scale follow-up imaging studies and the registration and correlation of the obtained data sets. Fig.4a) shows the 3D rendering of a PBI scan of a fixed mini-pig lung within the phantom. Following the scan, the lung was sliced as mentioned above and processed for histology. The obtained FFPE-tissue block was then scanned at 4.5 μm resolution. Fig.4b) shows the position of the FFPE-tissue block within the PBI CT data of the whole fixed lung. In Fig.4c), the PBI microCT data set is shown in greater detail. Clearly, major structures such as interlobular septa as well as the general distribution of alveoli, bronchi and vessels can be studied. In red, the location of a subsequently performed HRSRμCT scan (0.65 μm resolution) is indicated. Fig.4d) shows a 3D rendering of the HRSRμCT data set of a 2 mm punch out of the FFPE-tissue block. The presented data demonstrates that, based on the FA vapor fixation, the pig lung can be studied at multiple resolutions, spanning from the entire lung at 67 μm, to tissue level of 4.5 μm and even down to (sub)cellular resolution at 0.65 μm.

**Figure 4.**
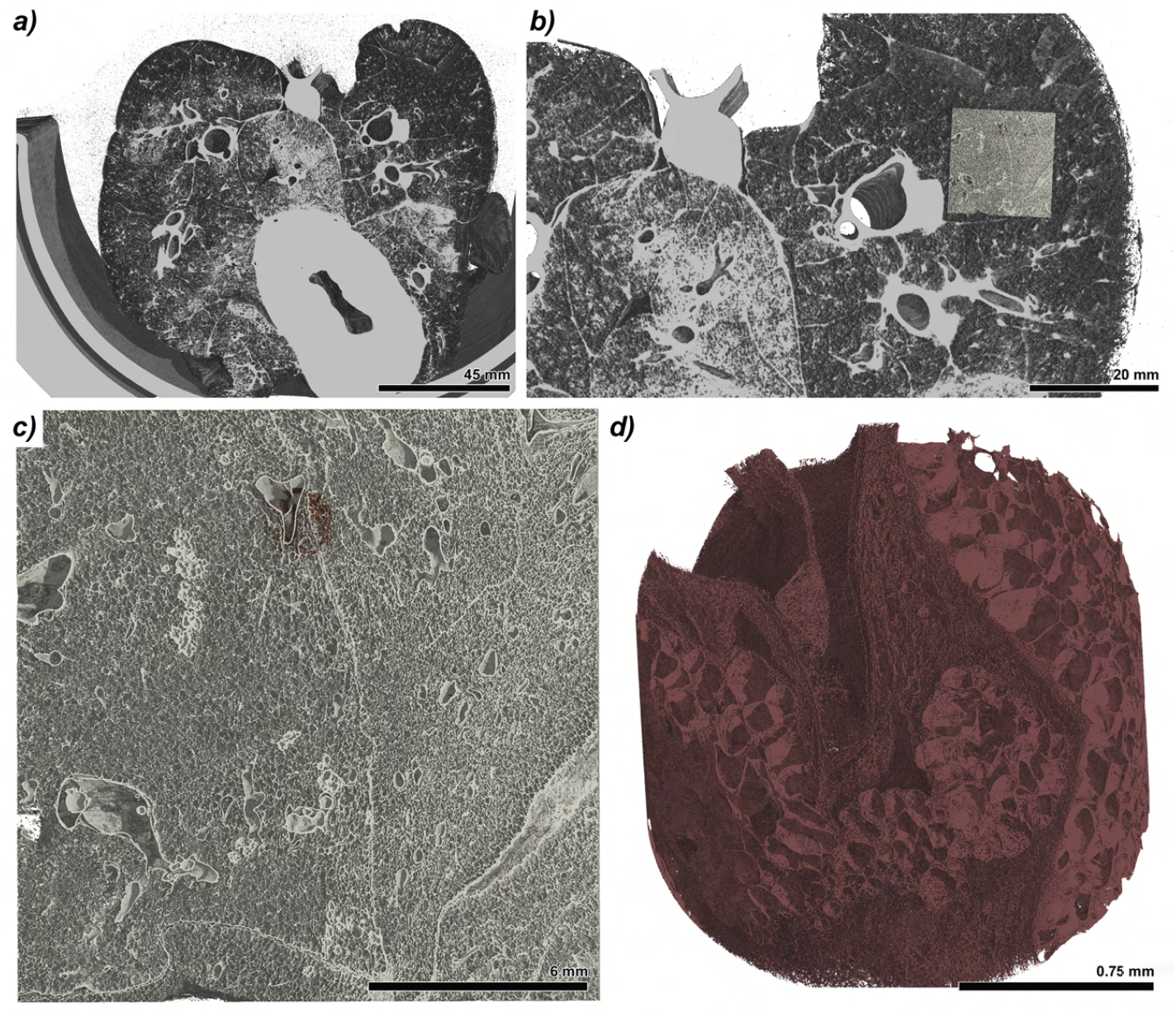
FA vapor fixation allows multi-scale imaging. (**a**) shows a 3D rendering of a FA vapor fixed lung scanned within the chest phantom at 67 μm resolution. Scale bar 45 mm. (**b**) shows a magnified region demonstrating the location of a FFPE-tissue block (yellow) after processing the lung. Scale bar 20 mm. (**c**) 3D rendering of a mosaic PBI acquisition of the entire paraffin block at 4.5 μm. In red, the position of a subsequently performed high resolution scan of 2 mm punch biopsy is marked. Scale bar 6 mm. (**d**) 3D rendering of a high resolution PBI scan of a tissue cylinder of the aforementioned FFPE-tissue block at 650 nm resolution. Scale bar, 0.75 mm.

### FA vapor fixed lungs show similar, if not better tissue quality in conventional histological imaging

Apart from the fact that the FA vapor fixation approach needs to be compatible with PBI scanning, it is also crucial that the tissue quality for subsequent histological examination is at least as good as for the classical approach of instilling FA into the trachea and immersing the entire lung into a FA bath. Fig.5a) shows an overview of a H&E stained slice from the top of the FFPE-tissue block of a FA vapor fixed mini-pig lung. The highlighted region demonstrates the location of the top slice of the retrospectively performed 2 mm diameter punch scanned with HRSRμCT at a resolution of 650 nm, which is displayed as 3D rendering in Fig.5b). Both the histological slide as well as the 3D rendered data confirm an overall well preserved tissue quality. For further validation, Fig.5c) and d) show detailed views with increasing magnification. For comparison, in Fig.5e) and f) H&E stained porcine lung tissue from a pig lung filled and immersed in FA are shown. Except for the fact that the expansion state of the lung tissue is different as evidenced by the differences in the shape of the alveoli (round shaped in Fig.5d) and partially crumbled in Fig.5f)), the overall quality of the FA vapor fixed specimen seemed to be at least comparable to the one prepared using the classical protocol of filling the airways of the lung with FA. To further elucidate this, we scored 4 features (size of the alveoli, homogeneity of the size distribution of the alveoli, thinness of the alveolar walls and absence of artifacts and ruptures). Each feature was scored between 0 and 3, with 0 = bad and 3 = very good. The scoring was performed by 4 independent readers, in a blind manner on 4 sections from each fixation technique. No significant differences in the four image features were found Suppl.Fig.S1.

**Figure 5.**
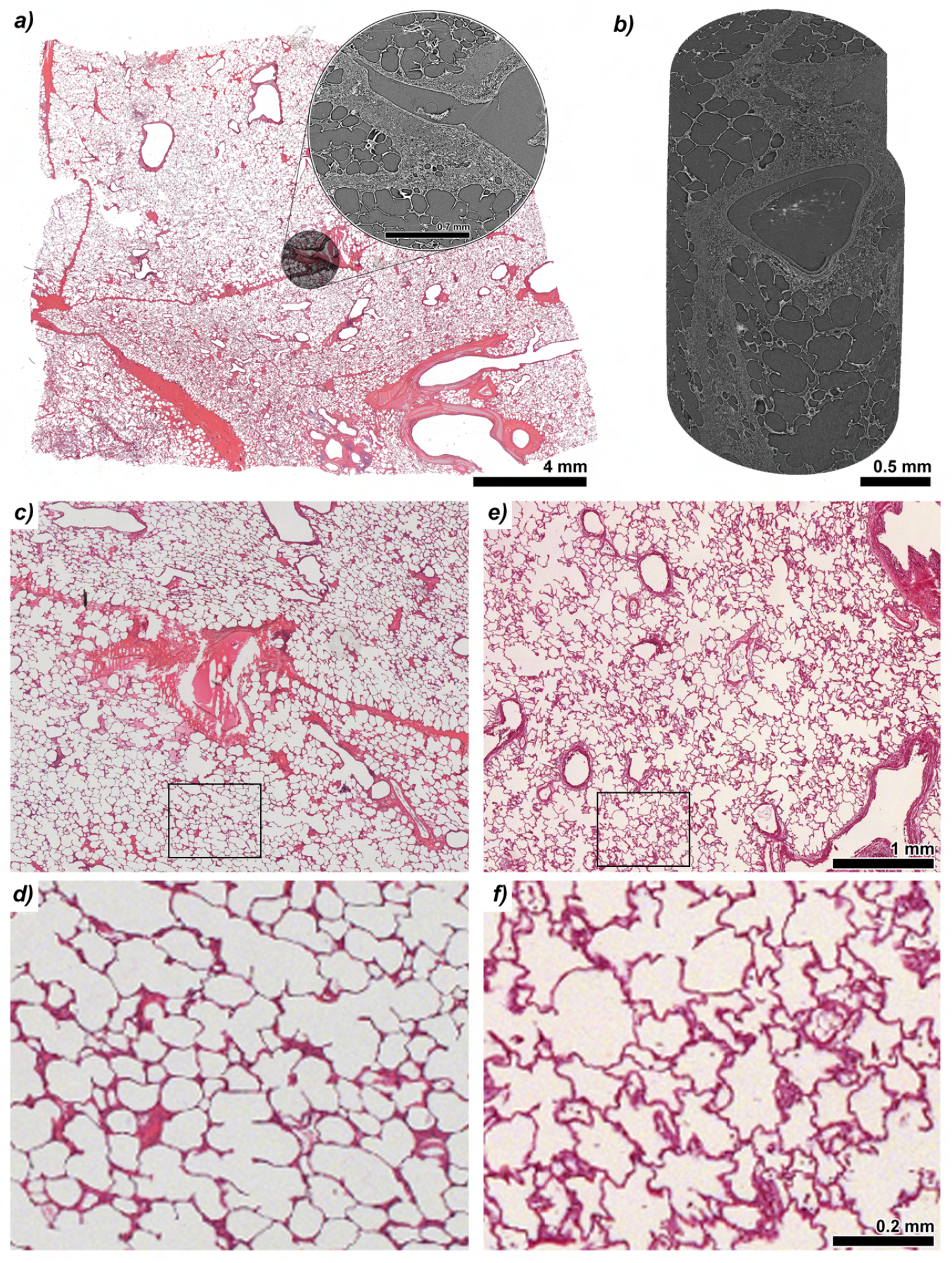
Comparison of histological quality between FA vapor fixation and lung tissue conventionally filled and immersed in FA. (**a**) shows a H&E stained slide from a FFPE-tissue block of a porcine lung fixed with FA vapor. In grey, the position and appearance of a high resolution PBI scan of a 2 mm diameter punch is overlaid. Scale bar 4 μm. Insert, Scale bar 0.7 mm. (**b**) 3D rendering of the 2 mm punch scanned at 650 nm resolution. Scale bar 0.5 mm. (**c**) magnified region of the same histology of the FA vapor fixed lung. (**d**) further magnified region indicated by the black rectangle in **c**. (**e**) histological section of a lung fixed by instilling FA and immersing the whole specimen in FA for multiple days. Scale bar 1 mm. (**f**) magnified region indicated by the black rectangle in **e**. Scale bar, 0.2 mm. Despite that the vapor fixed lung seems to be fixed in a more inflated state, no remarkable differences in the quality of the histologies were found.

### The workflow is compatible with clinical bronchoscopy

Since the phantom uses negative pressure between the lung and artificial chest wall to let the lung inflate passively via the trachea, the trachea remains accessible. This allows to perform bronchoscopy and to obtain biopsies inbetween the imaging sessions, close to the clinical diagnostic workflow in patients. In a pilot experiment we demonstrated that this can be done while the phantom is mounted at the PBI setup position. Suppl.FigS2a) shows the authors performing the bronchoscopy. Suppl.Fig.S2b) depicts a still frame of an acquired video and Suppl.Fig.S2c) demonstrated roughly the corresponding region in the retrospectively acquired PBI data set. Please note that in this first test the lung was ruptured and therefore could not be kept fully inflated during the PBI acquisition. Therefore, the CT data does not precisely match the bronchoscopy videos. The approach of combining the image acquisition with bronchoscopy examination seems feasible but needs further improvement. However, Suppl.Fig.S2d) and e) clearly demonstrate how strongly the structure of the obtained biopsies is compromised by the extraction process.

### Current limitations of the presented setup

It is crucial for the FA vapor fixation process to maintain the correct and consistent negative pressure to keep the lung inflated. As the lung tissue gets progressively stiffer in the process the suction rate of the pump might need to be adapted and monitored. Fig.6a) shows a slice through a PBI data set of a fresh mini-pig lung within the phantom which was scanned using the pressure pump. Fig.6b) shows a slice at approximately the same anatomical position in a PBI scan of the same lung after FA vapor fixation. Apparently, the negative pressure used during the fixation was too high and resulted in ruptures of the lung (**§**). In addition, the size of the lung was larger after the fixation compared to before (**yellow line**). Since the trachea of this specimen was short and damaged, the connection to the tube of the phantom was not completely sealed. Thus, not only the FA vapor but also regular air was sucked into the lung, which led to an incomplete fixation of the periphery of the lung as evidenced by the intensity gradient close to the surface (Fig.6b detail view lower right corner). When the FA vapor was cooled, too much liquid was entering the lung, obstructing the airways as seen in Fig.6c) (**\***). This again indicates the importance of the FA vapor fixation approach for PBI imaging in contrast to classical fixation. If the negative pressure during the fixation process is too low (Fig.6d) the lung tissue appears to be denser overall (in comparison to Fig. 6a) and the outer shape is irregular. Fig.6e) shows a lung specimen which was partially ruptured and not well inflated as well as liquid deposition areas (**bright regions**). Especially in the latter case, a characterization of pathological changes within the lung would be severely impaired by these artifacts. Thus, maintaining the correct and constant negative pressure and a steady stream of FA vapor without influx of liquid is crucial for the quality of the fixation process. Please note that the generation of FA vapor by boiling FA is also a difficult aspect. The FA concentration within the liquid increases over time which can result in a chemical precipitation of the salt used to bind the FA in the solution. To overcome these factors, the next step will be to utilize a fog machine that generates cold FA vapor.

**Figure 6.**
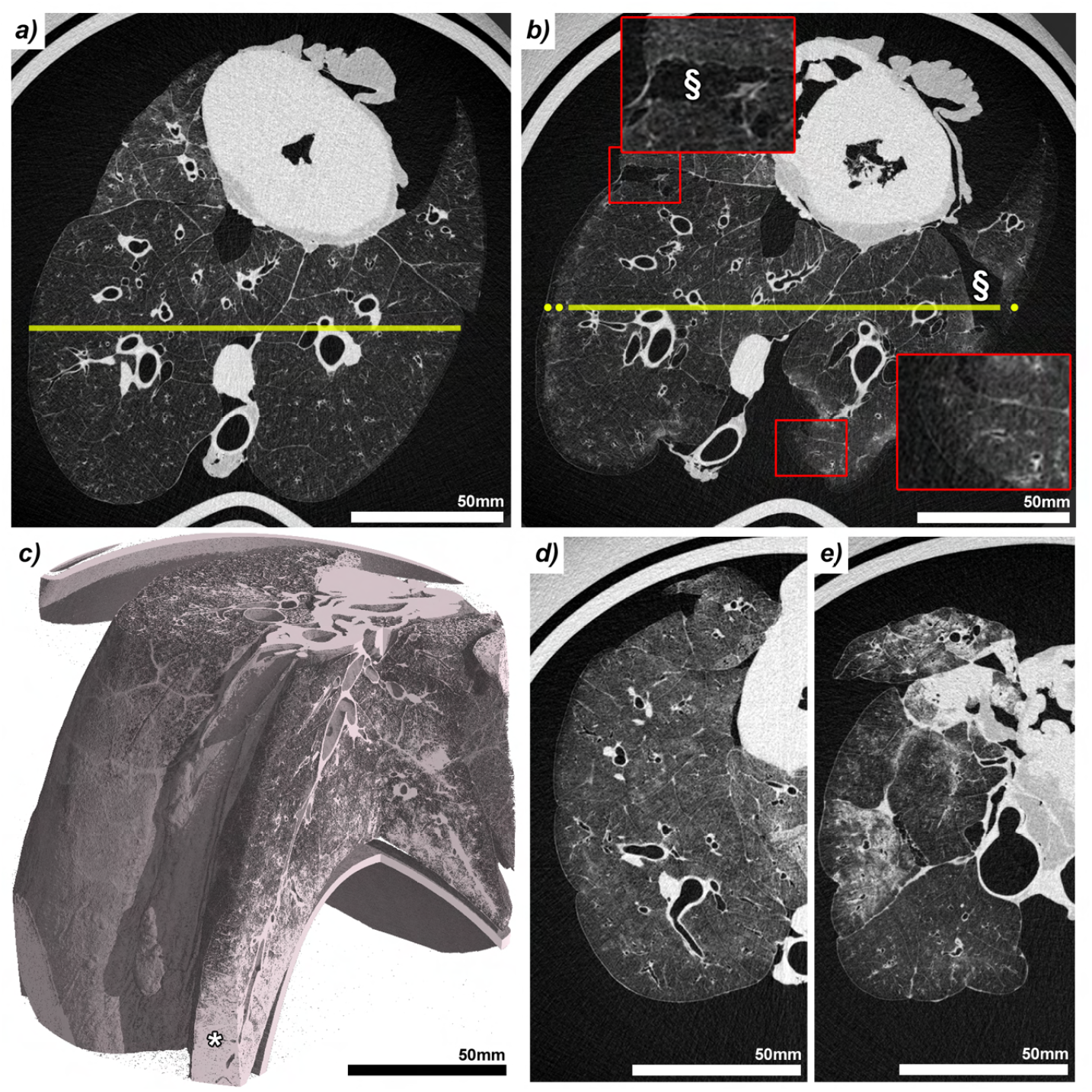
Problems in the FA vapor process. (**a**) and (**b**) show the same lung scanned within the phantom prior to fixation and using negative pressure **a** and after FA vapor fixation and without the application of negative pressure **b**. The entire lung volume has increased during fixation (yellow line) and apparent ruptures (§) were found, both pointing to an inadequately strong negative pressure during the fixation process. Moreover, the lung was very small and therefore it was difficult to fix the trachea to the pipe in the phantom, resulting in a leaky connection. Consequently, an insufficient amount of vapor entered the lung and the periphery of the specimen remained unfixed (second insert). Scale bar 50 mm. (**c**) shows a 3D rendering of a PBI scan of a lung in which not only FA vapor but also liquid FA entered the lung. The FA filled lung tissue does not provide sufficient contrast in PBI (asterisk), highlighting the importance of vapor fixation as opposed to instilling PFA or FA. (**d**) demonstrates a case in which the pressure during the fixation process was too low resulting in a more compact appearance and a deformed and overall smaller surface. (**e**) shows a case in which the lung was damaged and also liquid FA entered the organ during the fixation process, creating the regions with increased density which could be easily mistaken for pathological changes. Scale bars, 50 mm.

## Discussion

We presented an adapted FA vapor fixation technique based on the work of Weibel and Vidone (1961). We showed that this approach can be applied to porcine lungs mounted in an anthropomorphic human chest phantom (ANTHONY) designed to perform comparative imaging mimicking a human patient. Direct comparison of PBI scans of the same lungs prior and after FA vapor fixation revealed no differences and especially no deterioration in image quality. The fixed lungs became rigid and maintained the expansion state of the fixation process to a high degree. Consequently, the handling of the aforementioned samples was straight-forward, and the tissue specimens could be readily obtained. We further demonstrated that this approach allows subsequent multi-scale imaging and the validation of the PBI imaging by histological analysis. Furthermore, we showed that the design of the phantom allows to combine the PBI imaging with clinical bronchoscopy, which paves the way for comprehensive studies in larger lungs as close as possible to the clinical routine.

The human lung is a very complex organ in which only roughly 1/2 L of tissue enclose several liters of air Weibel (2009). Once removed from the body this delicate structure collapses, severely hindering quantitative analysis by histology, stereology or other methods. Furthermore, this also limits the ability to correlate detected pathological changes in in vivo high resolution CT with tissue specific methods. In our case, the performed PBI scans of the entire lung had a resolution of 67 μm which is much better than even the newly developed systems based on photon-counting detectors, which can reach an in-plane resolution of approximately 0.2 mm Remy-Jardin et al. (2023). In light of the development of these new imaging techniques, it is even more important to precisely validate the imaging findings. Knudsen et al. (2023) highlighted the importance of a controlled expansion state of the lung during fixation as a prerequisite for quantitative comparison. In our approach, since the flow-rates during the FA vapor fixation process change, more emphasis must be put on a precise pressure monitoring and regulation system, but we believe that our setup can be adapted in this perspective.

The FA vapor fixation process proposed by Weibel and Vidone (1961) has also been applied by King et al. (2000) to test high resolution CT. Their data shows the same bright areas that we observed whenever not only vapor but also liquid enters the lung, highlighting the importance of avoiding this effect.

Another critical aspect is the shrinkage of the lung after fixation. Weibel and Vidone (1961) reported that the vapor fixed lung still retained some ability to recoil and observed a shrinkage of 18% in linear dimensions. We also observed shrinkage in the vapor fixed lungs - however, the limited number of specimens and the only rough estimation of the amount of negative pressure and the suction rate does not allow a comprehensive analysis. Additional experiments are needed to study this effect. A slight and defined overinflation of the lung during the fixation process may compensate this effect, but would need to be applied carefully to avoid damage to the lung.

The initial pilot experiments conducted to develop high-resolution PBI lung CT in patients employed the use of the presented anthropomorphic human chest phantom. However, these experiments were constrained by the fact that only lungs of healthy pigs were used for the study. Pathologies were simulated using agarose gel mixed with iodine Albers et al. (2023). While this has proven to result in the typical radiological patterns of lung diseases, a histological validation could not be performed. Moreover, despite the fact the pig lung is similar in size to the human lung, there are also certain differences: i) it contains more cartilage which is found in more peripheral parts of the bronchial system and ii) the branching pattern is different to a human lung. Thus, it would be important to study pathologically altered human lung tissue. Naturally, fresh human lung tissue is difficult to be obtained. One potential avenue for obtaining this is a case of lung transplantation. However, such an approach is difficult to schedule. Since our setup is portable and only needs minimal setup time, human lung tissue could be FA vapor fixed quickly after explanation and subsequently analyzed with PBI and histology.

## Conclusion

We developed an adapted FA vapor fixation approach that enables a) the validation of high resolution CT imaging results in studying human sized lungs under similar conditions as in clinical CT by subsequent multiscale imaging and histology and b) the fixation of lungs in a way that, as subsequent PBI analysis reveals, provides the same image quality. This technique is of great value as it enables to study human lung tissue via PBI at clinical CT conditions as well as to validate the findings by subsequent histology. Thus, the approach will be fundamental for establishing novel clinical lung CT imaging techniques developed for the examination of human patients.

## Methods

### Specimens and sample preparation for CT scanning

Lungs of adult domestic pigs (N=5, body mass approximately 80 kg) have been obtained from a licensed slaughterhouse as a byproduct of food production. Two lungs of mini-pigs (body mass approximately 40 kg) have been obtained from the Animal Research Center (ARC) of the Technical University Munich (TUM) from a licensed sarcoma study in pigs (animal permit number: 55.2-2532.Vet_02-18-33). No animal was sacrificed for the purpose of this study.

Lungs were stored at −20°C for several days or weeks and transported frozen to the Italian synchrotron facility. Prior to imaging the lungs were slowly defrosted at room temperature for about 12 hrs to ensure the absence of ice crystals within the tissue. The specimens were then cleaned from additional tissue such as fat between the lung lobes as well as larger pieces of the diaphragm with attached muscle layers. The trachea was cut to an appropriate length to fit the specimen into the chest phantom. In case of visible damage to surface of the lung, these areas were sutured and glued using superglue.

One additional lung of an adult domestic pig was fixed by instilling FA into the trachea and immersing it into a bucket filled with formalin. This lung was not scanned with PBI.

### Propagation based phase contrast CT imaging using an anthropomorphic chest phantom

In order to mimic a potential patient lung CT acquisition, the lungs were placed in an anthropomorphic human chest phantom (ANTHONY=**ANT**hropomorphic p**H**antom f**O**r lu**N**g tomograph**Y**, MiMEDA GmbH, Germany). The phantom consist of an inner shell in which the lung is placed and connected to the outside via the trachea and a tube. The inner shell is sealed and a negative pressure is applied using a flow pump to let the lung inflate passively. The inner shell is then placed into an outer shell which can either be mounted at the synchrotron setup or scanned in a clinical CT. The outer shell comes in two versions, both mimicking the absorption of the chest wall of a patient but one with and the other without additional artificial bone structures. The phantom is made from acrylic glass, allowing to visually inspect the expansion state of the lung. The assembly of the phantom is shown in Suppl.Fig.S3.

After defrosting, the lungs were placed into the phantom and scanned using the flow pump. Following the PBI CT scans, the lungs were kept in the inner shell of the chest phantom and FA vapor fixed as described above. The fixed lung was then scanned again with the same settings to compare the effect of the fixation on the resulting image quality.

Propagation based phase contrast CT imaging (PBI) was performed at the SYRMEP beamline of the Italian synchrotron using a monochromatic x-ray beam of 40 keV (ΔE ≈ 50 eV) and a sample-to-detector distance of 10.7 m. We utilized the single photon counting detector with a pixel size of 100 × 100 μm^2^ (Hydra, Direct Conversion, Sweden). Given the divergence of the beam this resulted in an effective pixel size of 67 μm. We used an additional aluminium filter of 3.75 or 7.75 mm and set a minimal energy threshold of 24 keV for the detector. The beam dimension was 141 × 4.5 mm^2^. 360°off-center scans with 3600 angular spaced projections were obtained. The exposure time was 10 ms per projection, resulting in a total acquisition time per full rotation of 36 s. To cover a larger portion of the lung, up to 94 vertically overlapped scans with an increment of 1.6 mm were used and the resulting data was stitched using a custom made python script.

Thermoluminescence dosimeters were used to estimate the x-ray dose resulting in a dose of ≈43 mGy and ≈23 mGy for 3.75 mm and 7.75 mm filters respectively.

### Bronchoscopy examination

For bronchoscopy, we utilized a single-use flexible bronchoscope Ambu® aScope™5 Broncho HD 5.6/2 (Ambu A/S, Baltorpbakken 13, 2750 Ballerup, Denmark).

### Sample fixation and histology

After the final PBI examination, the fixed lungs were removed from the phantom and stored in sealed buckets without adding additional FA. After 2 to 4 weeks, the lungs were removed from the bucket and sliced into approximately 5 mm thick slices while trying to cut at an angle parallel to the orientation of the reconstructed CT data. From these slices, pieces of about 2 × 3 cm^2^ were cut, chemically dehydrated using an ascending ethanol series and embedded in paraffin using the standard protocol of the pathology department. The whole slicing and embedding process was carefully documented to ease the subsequent localisation of the histology data within the PBI scans.

From the resulting formalin-fixed and paraffin-embedded (FFPE) lung tissue, 2 μm thick sections were obtained. We used the first section close to the surface of the paraffin block that depicted the entire specimen. Sections were then stained with Haematoxylin Eosin (H&E) using the established standard staining protocol. Mosaic microscopy images were obtained at 5 fold magnification to cover the entire histological specimen and details were recorded with 20 fold magnification both using an Axiovert 200 inverted microscope (Zeiss, Germany).

### PBI microCT examination

The whole paraffin embedded samples were scanned at the SYRMEP beamline using the white beam setup with a sample-to-detector distance of 500 mm. A 16-bit, water-cooled sCMOS camera (Hamamatsu C11440-22C ORCA-Flash 4.0 v2) was used to acquire 3600 angular distributed projections with an exposure time of 50 ms resulting in an isotropic voxel size of 4.5 μm. Scans were performed in a 360° offset regime, resulting in a scan time of 180 s. A 0.5 mm Si filter was used, resulting in a mean beam energy of 19.6 keV. To cover the main tissue area close to the surface of the paraffin block, a mosaic acquisition with either 2 × 3 or 2 × 4 tiles was performed.

### HRSRμCT examination

2 mm punch biopsies were scanned at the high throughput tomography (HiTT) setup at the EMBL P14 beamline on PETRA III (Hamburg, Germany) Albers et al. (2024). An X-ray microscope with a 10x objective coupled to a PCO.edge 4.2 sCMOS camera was used, resulting in an effective pixel-size of 0.650 μm. A monochromatic X-ray beam with a photon energy of 18 keV was used to acquire 3600 projection images over a rotation angle of 360°with an offset rotation axis to increase the lateral field of view (FOV). Four acquisitions per sample were performed at 73, 77, 83 and 92 mm sample-to-detector distance (SDD) respectively to enable phase retrieval with a contrast-transfer-function (CTF) based algorithm. An exposure time of 10 ms per projection was used resulting in a total exposure time of 144 s per acquisition. For each specimen three adjacent scans with a vertical offset of 0.9 mm were acquired resulting in a FOV of approximately 2.6 × 2.6 × 3.1 mm^3^. Phase retrieval and reconstructions was performed using the in-house made TOMO-CTF software. A δ/β ratio of 10 was used. Reconstructions were stitched together using NRStitcher.

### Software and statistics

All data sets acquired at the SYRMEP beamline were reconstructed using the SYRMEP Tomo Project software (STP Version 1.6.3) Brun et al. (2015). A single distance phase retrieval algorithm developed by Paganin et al. Paganin et al. (2002) was used with a *δ/β* ratio of 2000 for the whole lung and 100 for the FFPE-tissue scans. Data was preprocessed in Fiji (NIH) Schindelin et al. (2012). For stiching of the mosaic white-beam acquisition NRStitcher Miettinen et al. (2019) was applied. Vertical stitching of the whole lung PBI scans was achieved with a custom made python script based on scikit-image phase cross-correlation function Guizar-Sicairos et al. (2008). Statistical analysis was performed using the python packages seaborn Waskom (2021) and statanot (https://pypi.org/project/statannot/). Rendering of the 3D data sets was performed with VGStudio Max v3.1 (Volumegraphics, Germany). Figures were generated with Photoshop 6.0 (Adobe, USA).

## Additional Information

### Ethics

No animal was sacrificed for the particular purpose of this study. Two lungs were obtained from an approved experiment using a sarcoma mini-pig model (animal permit number: 55.2-2532.Vet_02-18-33) after its completion. All other lungs were obtained from a licensed slaughterhouse. The preparations passed the regular veterinarian controls of a licensed slaughterhouse and were treated with the same hygiene precautions as fresh meat. Storage, transport, handling and disposal of faunal by-products were registered at the responsible veterinary office. All methods were carried out according to national law (German law). The pigs were sacrificed with the appropriate euthanasia methods, using a captive bolt pistol in case of the pigs from the slaughterhouse. In case of the pigs from the sarcoma model, the pigs were sedated by IM administration of ketamine (20 mg/kg body weight) and azaperone (2 mg/kg body weight), rendered unconscious by a captive bolt gun applied to the forehead, and then exsanguinated.

## Acknowledgements

The authors acknowledge the support in tissue embedding, cutting, staining and microscopy by Sarah Garbode and Stefan Lesemann (University Medical Center Goettingen) as well as Jolanthe Schatterny (Translational Lung Research Center, Heidelberg). We also thank Hermann Nolte for his help in organizing the lung specimens.

## Author contributions

C.D., J.R., W.W. and G.T. designed the experiment. C.D., J.R., E.L., M.P., A.C., P.C., E.B., J.A., A.S., T.F., C.F., F.Z., C.B. and G.T. performed the experiment. C.D., J.R., S.M., J.A., A.S., M.S. analyzed the data. All authors contributed to the writing process and have approved the final manuscript.

## Supplementary Information

**Figure S1.**
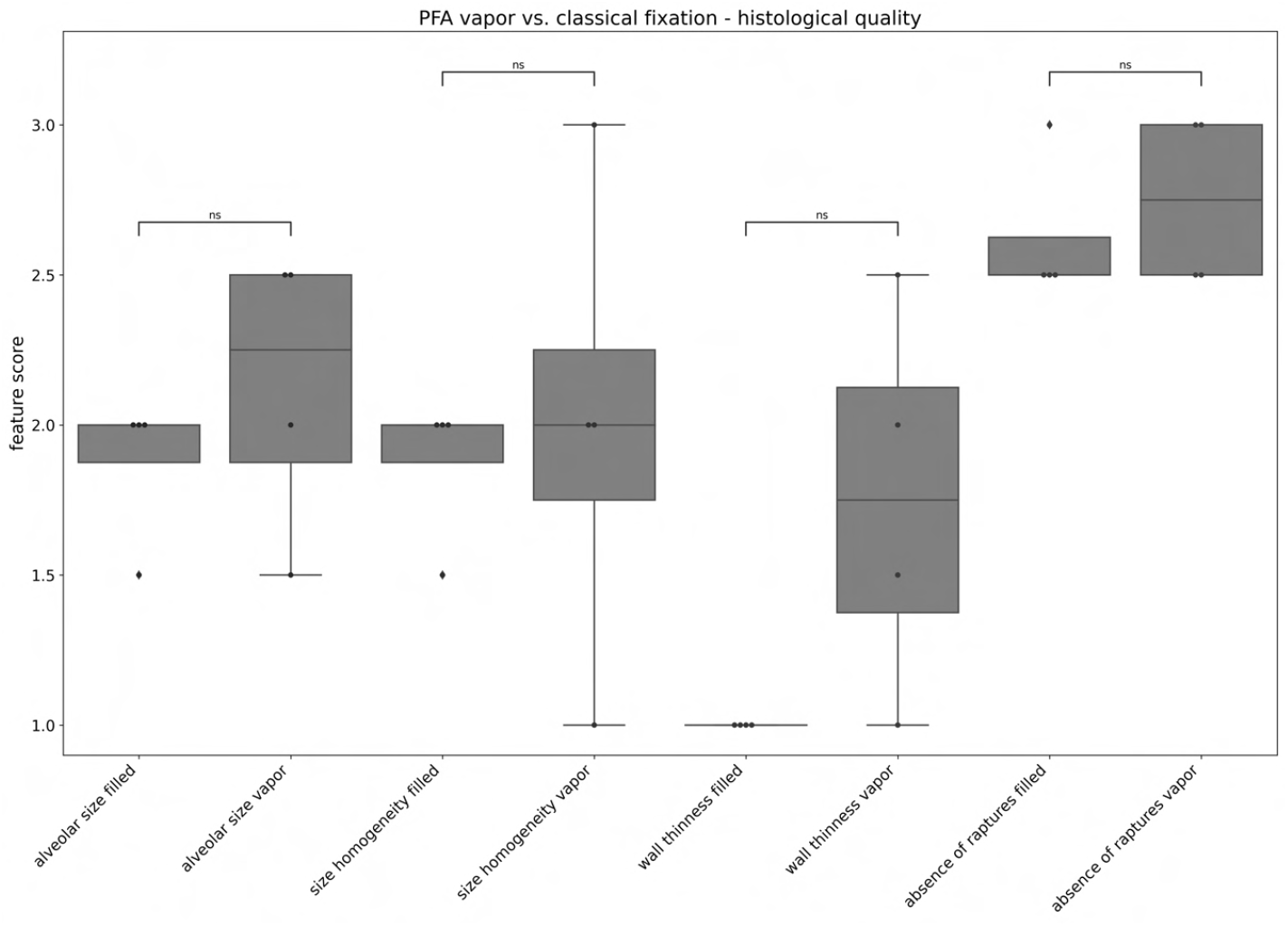
Scoring of histological image quality. Four different features (**alveolar size**,**homogeneity of alveolar sizes, thinness of alveolar walls** and **absence of raptures**) was scored between 0 (bad) and 3 (very good) by 4 independent readers in a blind and randomized manner. The data is shown comparing classical fixation by instilling PFA to the trachea (**filled**) and by the presented vapor fixation approach (**vapor**). While the vapor fixation was in average scored slightly higher than the classical approach, none of the differences were found significant.

**Figure S2.**
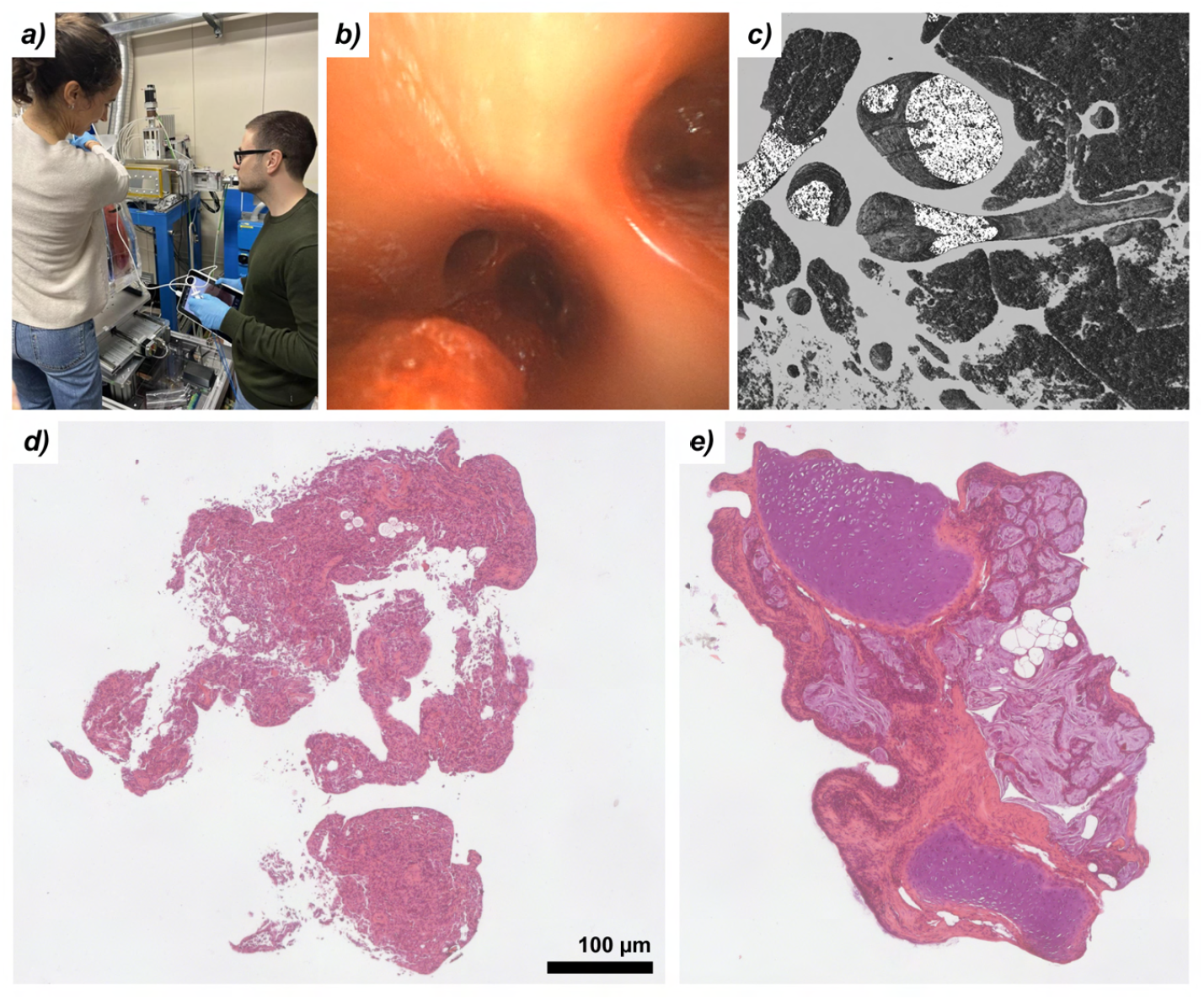
Combining PBI and bronchoscopy. (**a**) due to the fact that the lung is passively inflated the trachea remains open and accessible. In between the imaging session bronchoscopy can be performed as shown here for the phantom mounted at the Synchrotron setup. (**b**) depicts a still frame of the obtained bronchoscopy video. (**c**) 3D rendering of the PBI data set of approximately the region shown in **b**. Please note that in this pilot study the lung was damaged between bronchoscopy examination and PBI scan resulting in deformation of the structure. (**d** and **e**) show H&E stained slices of two of the obtained biopsies. Clearly the structure of the tissue was strongly compromised by the extraction process.

**Figure S3.**
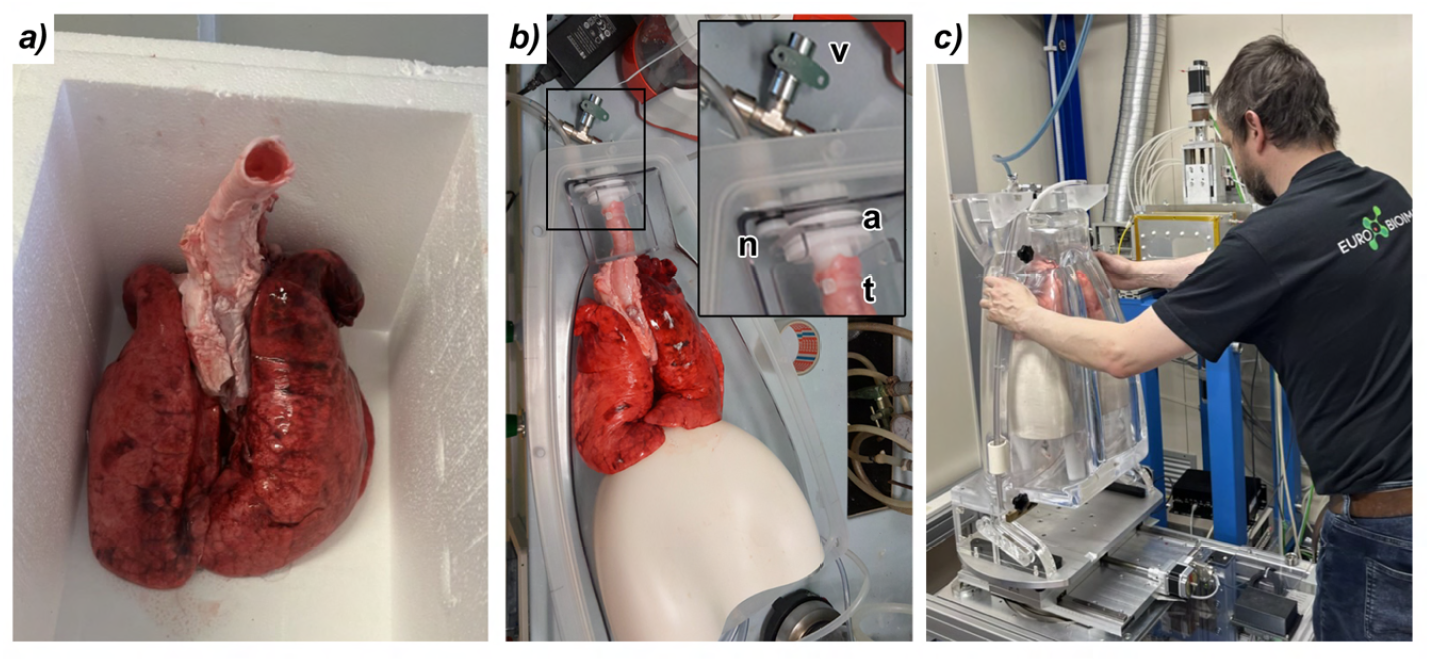
Phantom design and placement of the lung specimen. (**a**) pig lung at the end of the defrosting process. Needles tissue was already removed and the trachea was cut to an appropriate length. (**b**) Lung placed in the inner shell of the phantom. Insert shows the drilling **n** to apply the negative pressure within the phantom. The three-way valve **v** can be used to adjust the sucking rate of the pump. The trachea **t** is fixed with a cable strap to the pipe that provides access to the surrounding air. (**c**) the author placing the inner shell with the equipped lung into the outer absorber shell mounted to the synchrotron setup.

## Bibliography

Albers, J., Wagner, W. L., Fiedler, M. O., Rothermel, A., Wünnemann, F., Di Lillo, F., Dreossi, D., Sodini, N., Baratella, E., Confalonieri, M., et al. High resolution propagation-based lung imaging at clinically relevant x-ray dose levels. Scientific Reports, 13(1):4788, 2023.

Albers, J., Nikolova, M., Svetlove, A., Darif, N., Lawson, M. J., Schneider, T. R., Schwab, Y., Bourenkov, G., and Duke, E. High throughput tomography (hitt) on embl beamline p14 on petra iii. Journal of Synchrotron Radiation, 31(1), 2024.

Brun, F., Pacilè, S., Accardo, A., Kourousias, G., Dreossi, D., Mancini, L., Tromba, G., Pugliese, R., et al. Enhanced and flexible software tools for x-ray computed tomography at the italian synchrotron radiation facility elettra. Fundamenta Informaticae, 141 (2-3):233–243, 2015.

Guizar-Sicairos, M., Thurman, S. T., and Fienup, J. R. Efficient subpixel image registration algorithms. Optics letters, 33(2):156–158, 2008.

King, G. G., Muller, N. L., Whittall, K. P., Xiang, Q.-S., and Pare, P. D. An analysis algorithm for measuring airway lumen and wall areas from high-resolution computed tomographic data. American journal of respiratory and critical care medicine, 161(2):574–580, 2000.

Kitchen, M. J., Buckley, G. A., Gureyev, T. E., Wallace, M. J., Andres-Thio, N., Uesugi, K., Yagi, N., and Hooper, S. B. Ct dose reduction factors in the thousands using x-ray phase contrast. Scientific reports, 7(1):15953, 2017.

Knudsen, L., Hummel, B., Wrede, C., Zimmermann, R., Perlman, C. E., and Smith, B. J. Acinar micromechanics in health and lung injury: what we have learned from quantitative morphology. Frontiers in Physiology, 14:1142221, 2023.

Miettinen, A., Oikonomidis, I. V., Bonnin, A., and Stampanoni, M. Nrstitcher: non-rigid stitching of terapixel-scale volumetric images. Bioinformatics, 35(24):5290–5297, 2019.

Paganin, D., Mayo, S. C., Gureyev, T. E., Miller, P. R., and Wilkins, S. W. Simultaneous phase and amplitude extraction from a single defocused image of a homogeneous object. Journal of microscopy, 206(1):33–40, 2002.

Remy-Jardin, M., Hutt, A., Flohr, T., Faivre, J.-B., Felloni, P., Khung, S., and Remy, J. Ultra-high-resolution photon-counting ct imaging of the chest: a new era for morphology and function. Investigative Radiology, 58(7):482–487, 2023.

Schindelin, J., Arganda-Carreras, I., Frise, E., Kaynig, V., Longair, M., Pietzsch, T., Preibisch, S., Rueden, C., Saalfeld, S., Schmid, B., et al. Fiji: an open-source platform for biological-image analysis. Nature methods, 9(7):676–682, 2012.

Waskom, M. L. seaborn: statistical data visualization. Journal of Open Source Software, 6 (60):3021, 2021. doi: 10.21105/joss.03021.

Weibel, E. E. and Vidone, R. A. Fixation of the lung by formalin steam in a controlled state of air inflation. American Review of Respiratory Disease, 84(6):856–861, 1961.

Weibel, E. R. What makes a good lung? Swiss medical weekly, 139(2728):375–386, 2009.

